# Asparagine synthetase and G-protein coupled estrogen receptor are critical responders to nutrient supply in *KRAS* mutant colorectal cancer

**DOI:** 10.1101/2023.05.05.539577

**Authors:** Lingeng Lu, Qian Zhang, Xinyi Shen, Pinyi Zhen, Audrey Marin, Rolando Garcia- Milian, Jatin Roper, Sajid A. Khan, Caroline H. Johnson

## Abstract

The nutrient status of the tumor microenvironment has major impacts on cell growth. Under nutrient depletion, asparagine synthetase (ASNS)-mediated asparagine production increases to sustain cell survival. G protein-coupled estrogen receptor-1 (GPER1) signaling converges via cAMP/PI3K/AKT with KRAS signaling to regulate *ASNS* expression. However, the role of GPER1 in CRC progression is still debated, and the effect of nutrient supply on both *ASNS* and *GPER1* relative to *KRAS* genotype is not well understood. Here, we modeled a restricted nutrient supply by eliminating glutamine from growing cancer cells in a 3D spheroid model of human female SW48 *KRAS* wild-type (WT) and *KRAS* G12A mutant (MT) CRC cells, to examine effects on *ASNS* and *GPER1* expression. Glutamine depletion significantly inhibited cell growth in both *KRAS* MT and WT cells; however, *ASNS* and *GPER1* were upregulated in *KRAS* MT compared to WT cells. When nutrient supply was adequate, *ASNS* and *GPER1* were not altered between cell lines. The impact of estradiol, a ligand for GPER1, was examined for any additional effects on cell growth. Under glutamine deplete conditions, estradiol decreased the growth of *KRAS* WT cells but had no effect on *KRAS* MT cells; estradiol had no additive or diminutive effect on the upregulation of *ASNS* or *GPER1* between the cell lines. We further examined the association of *GPER1* and *ASNS* levels with overall survival in a clinical colon cancer cohort of The Cancer Genome Atlas. Both high *GPER1* and *ASNS* expression associated with poorer overall survival for females only in advanced stage tumors. These findings suggest that *KRAS* MT cells have mechanisms in place that respond to decreased nutrient supply, typically observed in advanced tumors, by increasing the expression of *ASNS* and *GPER1* to drive cell growth. Furthermore, *KRAS* MT cells are resistant to the protective effects of estradiol under nutrient deplete conditions. ASNS and GPER1 may therefore be potential therapeutic targets that can be exploited to manage and control *KRAS* MT CRC.

## Introduction

Metabolic reprogramming is a hallmark of various human cancers including colorectal cancer (CRC), where it facilitates tumor growth by adapting to insufficient nutrient supply and hypoxia (1). In addition to the Warburg effect, which preferentially increases the rate of glucose uptake and production of lactate for energy production over oxidative phosphorylation, it is known that cancers are also addicted to glutamine. This suggests that as a non-essential amino acid, glutamine has been switched to an essential amino acid in cancer; without exogenous glutamine, cancer cells cannot survive and grow (2). As a substrate, glutamine provides obligate nitrogen for the synthesis of nucleotides (purine and pyrimidine synthesis) and non-essential amino acids. Notably, the non-essential amino acid asparagine, which is catalyzed by asparagine synthetase (ASNS), has been recognized as an essential amino acid in cancer, supporting cell growth under stress and nutrient poor conditions (3). Asparagine can rescue glutamine depletion-induced cell inhibition (4), and in response to nutrient depletion, ASNS-mediated asparagine production is increased to sustain cancer cell survival (5).

Somatic *KRAS* mutations are found in 40% of CRCs (6). Via guanine nucleotide exchange factors (GEF), GTP is bound to KRAS, replacing GDP, which switches on KRAS signaling. In contrast, KRAS signaling is switched off by GTPase-activating proteins (GAP), which catalyze the hydrolysis of KRAS-GTP. Mutations can alter the conformation of the KRAS protein that block the binding of GAP to KRAS. This leads to constitutive KRAS signaling, and the activation of downstream molecules that are involved in biological processes such as cell proliferation, migration, angiogenesis, and DNA synthesis (7). In addition, the activation of downstream molecules such as activating transcription factor 4 (ATF4) upregulate the expression of ASNS (7, 8). Consequently, the increase of intracellular asparagine, due to the activation of the KRAS signaling pathway, further stimulates cell proliferation, activates mammalian target of rapamycin complex 1 (mTORC1), but suppresses apoptosis, thereby promoting tumor development and progression (7).

Reduction of intracellular asparagine by different strategies is an approach to effectively control tumor metastasis of human cancer, e.g., breast and pancreatic cancer, in preclinical studies (7, 9, 10). For example, L-asparaginase has been approved by the US Food and Drug Administration for the treatment of acute lymphoblastic leukemia (ALL), which converts asparagine into aspartate, consequently lowering the concentration of circulating asparagine. However, it has been reported that the effect of decreasing asparagine levels could result in upregulation of *ASNS* expression in a feedback loop. This would stimulate asparagine production, consequently resulting in resistance to the anti-cancer activity of L-asparaginase that has been reported in solid human cancers (11). Furthermore, the presence of *KRAS* mutations could adapt CRC cells to conditions of glutamine deficiency by increasing *ASNS* and consequently asparagine biosynthesis; *ASNS* knockdown has led to the growth suppression of *KRAS* mutant CRC *in vivo* (4). It has also been proposed that mutant KRAS can promote glutaminolysis and *ASNS* expression, activate WNT signaling, and increase metastasis (12). In addition, we recently reported that there was a significant positive association between high asparagine levels, high *ASNS* expression, and poor prognosis in female patients with CRC. This association was not seen in males, suggesting that ASNS and asparagine may have sex-specific effects on CRC progression and prognosis (13, 14).

The hormone estrogen may play a protective role in the development and progression of CRC. An approximate 56% CRC risk reduction was reported in postmenopausal women given hormone replacement therapy (HRT) in the Women’s Health Initiative (WHI) trial (15). The Prostate, Lung, Colorectal and Ovarian (PLCO) cancer screening randomized trial also showed that current HRT users had a significantly reduced mortality risk compared to never-users (16). Women who used the oral contraceptive pill of either estrogen alone or in combination with progesterone had a reduced CRC risk of 30% (17). The protective effect of estrogen is thought to be through its activation of estrogen receptor beta (ERβ) which causes pro-apoptotic signaling, inhibition of inflammatory signals, and modulation of the tumor microenvironment (15). However, a recently discovered transmembrane estrogen receptor, G-protein coupled estrogen receptor (GPER1), is highly sensitive to the nutrient supply of the CRC microenvironment. Under normoxia, GPER1 agonists suppress cell proliferation and growth, while under hypoxia, GPER1 enhances cell growth (18). This difference may be attributed to the opposing actions of estrogen-GPER1 signaling on HIF1α and VEGFA, which stimulate CRC cell proliferation and inhibit cell apoptosis (19). More importantly, studies have shown links between GPER signaling and downstream molecules of KRAS (20–22). GPER1 signaling activation increases the activity of phosphoinositide 3-kinase (PI3K)/protein kinase B (AKT) via cyclic adenosine monophosphate (cAMP) (23–25). PI3K/AKT is a downstream target of KRAS which regulates the expression of *ASNS* via NRF2 (as known as NFE2L2) and ATF4 (7). However, the effect of GPER1 activation on CRC cell growth is still uncertain, and it is not known if it has any indirect effects on ASNS (26, 27).

These observations suggest that the sex-steroid hormone estrogen may regulate the effect of glutamine and asparagine on *KRAS*-mutant (MT) CRC in a different mechanism compared to *KRAS* wild-type (WT) CRC. However, most of these observations have been previously derived from 2D cell culture conditions. Thus, the purpose of this study is to investigate the effect of nutrient supply and the sex-steroid hormone 17β-estradiol on *GPER*, *ASNS* and cell growth in CRC. We use female *KRAS* WT and isogenic *KRAS* MT SW48 cells in an *in vitro* 3D organoid model, which better mimics in *vivo* hypoxia and nutrient depleted microenvironments in tumors than 2D cell culture models (28), and examine clinical data to determine the association between *GPER1* expression and patient survival in CRC.

## Materials and methods

### Cell lines and chemicals

SW48 *KRAS* G12A MT and its isogenic parent SW48 *KRAS* WT cell lines were purchased from Horizon Discovery Ltd (Cambridge, UK). Both cell lines were maintained in Gibco^TM^ complete phenol red-free high glucose Dulbecco’s Modified Eagle Medium (DMEM) (ThermoFisher Scientific inc., Waltham, MA) with 4 mM Gibco^TM^ glutamine or specific concentrations of glutamine as mentioned (ThermoFisher Scientific inc), and supplemented with 10% Gibco^TM^ heat-inactivated fetal bovine serum (FBS) and 1% Gibco^TM^ Penicillin-Streptomycin (10,000 U/mL), in a humidified incubator with 5% CO_2_ at 37°C. The medium was replaced every other day, and the cells were passaged routinely when 80-90% confluence was reached. The cell line KRAS mutation type was validated with RNA sequencing. 17β-Estradiol was purchased from Sigma-Aldrich (St Louis, MO). The concentration of the stock solution was 100 mM for estradiol in dimethyl sulfoxide (DMSO).

## 3D spheroid formation

3D organoid formation assay was performed using a hanging-drop approach. Each drop (20 µl) of freshly prepared SW48 *KRAS* WT or MT cell suspension (approximately 500 cells per drop) was applied. The cells were treated in the presence or absence of estradiol at a final concentration of 10 µM, and in the presence (4 mM) or absence of glutamine, the same volume of DMSO was used as a negative control. After 5 days of incubation with 5% CO_2_ at 37°C, images were captured using a Nikon Eclipse E200 (Nikon, Japan)-Axiocam system, which is supported by Zen^©^3.3 (Carl Zeiss Microscopy GmbH) software. The area and intensity of 3D spheres were determined using an embedded plugin ImageJ for each organoid. The size of the 3D spheres was calculated by multiplying the area and intensity. The inhibition rate was calculated using the formula of (1– ratio of 3D sphere size (condition A vs. B)×100% (e.g., glutamine deplete vs. replete). The cells were then harvested and re-suspended in the QIAzol lysis reagent (Qiagen, Germantown, MD) for total RNA extraction and next generation RNA-sequencing. A final concentration of 0.1% DMSO in the medium was used as a negative control.

### RNA sequencing and pathway analysis

Total RNA was extracted from the harvested cells on day 6 after the incubation in the QIAzol lysis reagent using miRNeasy mini kit (Qiagen) following the manufacturer’s instructions. The concentration and quality of total RNA were determined using Agilent analyzer (Santa Clara, CA). Poly(A) RNA libraries were made and paired-end (2× 100 bp) next generation RNA sequencing was conducted on an Illumina platform of NovaSeq S2 sequencer system (San Diego, CA) at Yale Keck Center for Genome Analysis (YCGA). RNA-sequencing data analysis was performed using Partek Flow software (Build version 10.0.23.0414, Partek, Inc., St. Louis, MO, USA). Reads were aligned to the genome (UCSC hg19) using STAR – 2.7.3a (29) and quantified to the transcriptome using Quantify to annotation model (Partek E/M) algorithm using Ensembl release 100. Differential expression was performed using the Bioconductor DESeq2 package (30). Results were filtered by a log2 fold change > 1 or < –1 and Benjamini–Hochberg False Discovery Rate (FDR) < 0.05 before performing pathway analysis (31). Functional pathway analysis was conducted using Ingenuity pathway analysis (IPA) (Qiagen, Redwood City, CA, Version 90348151).

### Colony formation assay

Freshly prepared cells were seeded in a six-well plate with approximately 500 cells/well. Cells were incubated in a 5% CO_2_ incubator at 37°C for 10 days, and the media were replaced with fresh media every other day. The cells were then fixed with 4% paraformaldehyde for 30 min and dyed with 0.2% crystal violet. The number of colonies were counted. The experiment was repeated in triplicate.

### GPER1 expression and clinicopathological data from a TCGA CRC dataset

Normalized *GPER1* mRNA levels and clinicopathological data were retrieved from a publicly accessible Genomic Data Commons (GDC) TCGA colon cancer (COAD) dataset (https://xena.ucsc.edu) using R package UCSCxenaTools. Patients with primary colon cancer tissues available for RNA-sequencing were included in this study. Binary *GPER1* expression level, high or low, was classified based on the cutoff value of *GPER1* expression level determined by an algorithm of the maximization of hazard ratio using R package survminer. Binary ASNS expression level, high or low, was classified based on the median as the cutoff value, to balance the sample size for subgroups. Overall survival in months was calculated based on the diagnosis date and either death or the last follow-up date, whichever came the first.

## Statistical analysis

Statistical analysis was conducted using R (version 4.0.5). Generalized linear model (GLM) or Wilcoxon rank sum test were appropriately used to assess the differences for continuous variables between groups with post-hoc Tukey’s test for multiple comparison correction. R package DESeq2 was used for differential expression of gene analysis with FDR multiple comparison correction. The average and variation of continuous variables were presented in mean (or median) and standard deviation (or range). Survival analysis in R package survival was performed using Kaplan-Meier survival curve with log-rank test. P < 0.05 was considered statistically significant.

## Results

### Glutamine depletion suppressed cell colony and 3D spheroid formation

Colony formation assays showed that decreasing the concentrations of glutamine from 4 mM to 0 mM significantly decreased cell colonies for both SW48 *KRAS* MT and WT cell lines (**Figures 1A–B**). Using a hanging-drop approach, both *KRAS* MT and WT cell lines were grown in either glutamine replete (4 mM) or deplete (0 mM) DMEM medium. We observed that the size of 3D organoids for both the *KRAS* MT and WT significantly decreased when grown in the glutamine-deplete medium in comparison to the 4 mM glutamine-replete medium (**Figures 2A–B**) (p < 0.001). On average, the inhibition rates were 74.5% for the *KRAS* MT, and 79.9% for the *KRAS* WT (**Figures 2A–B**). The inhibition rate of glutamine depletion was significantly higher for *KRAS* WT compared to *KRAS* MT cells (p = 9.62 × 10^-9^).

**Figure 1.**
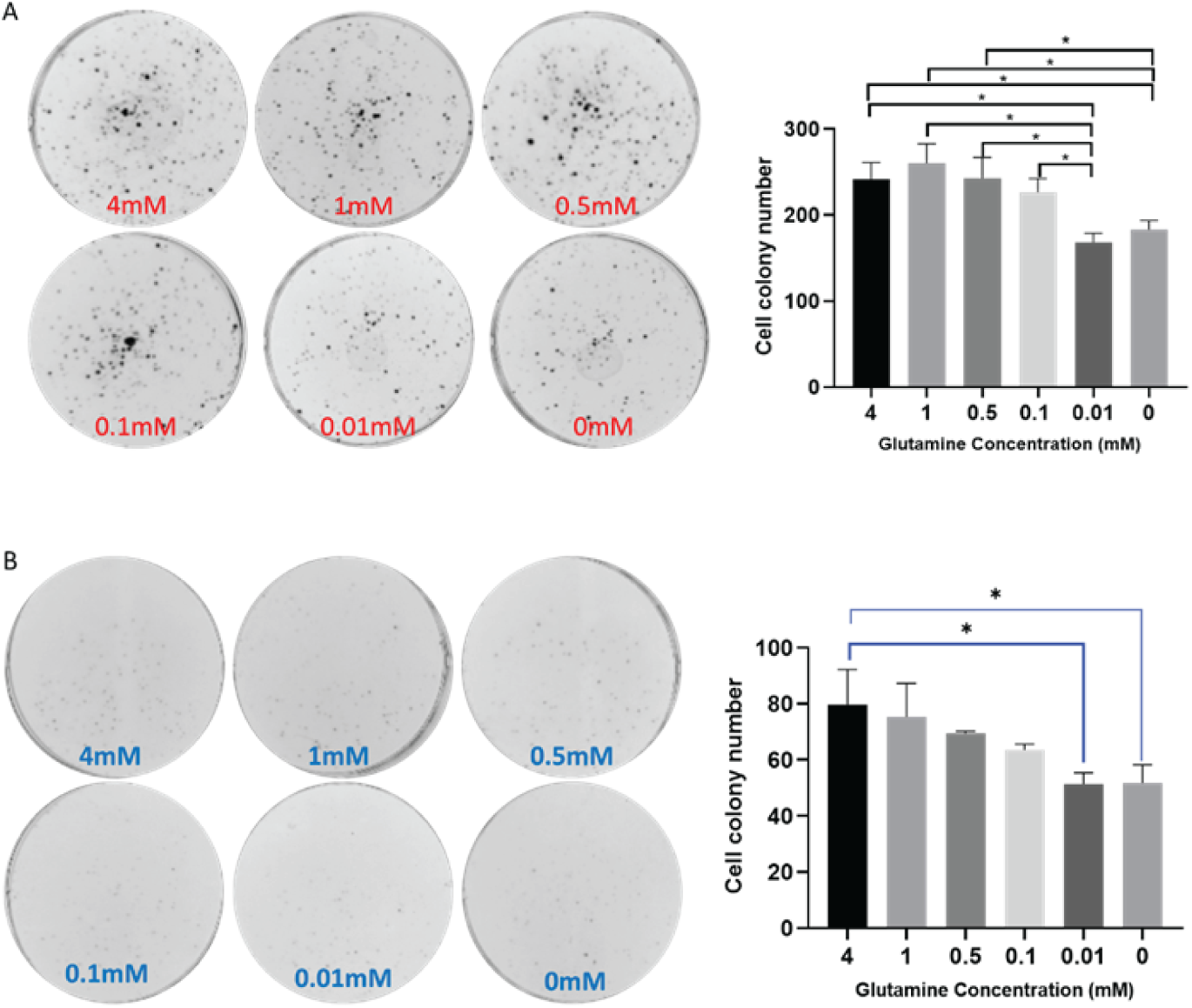
The inhibition of CRC SW48 cell colony formation *in vitro* under glutamine deplete conditions. (**A**) CRC SW48 *KRAS* MT and (**B**) CRC SW48 *KRAS* WT. Left panels are representative images of cell colony formation assays, and the right panels are quantitative data representing mean and standard deviation of six replicates under each condition * = p<0.05.

**Figure 2.**
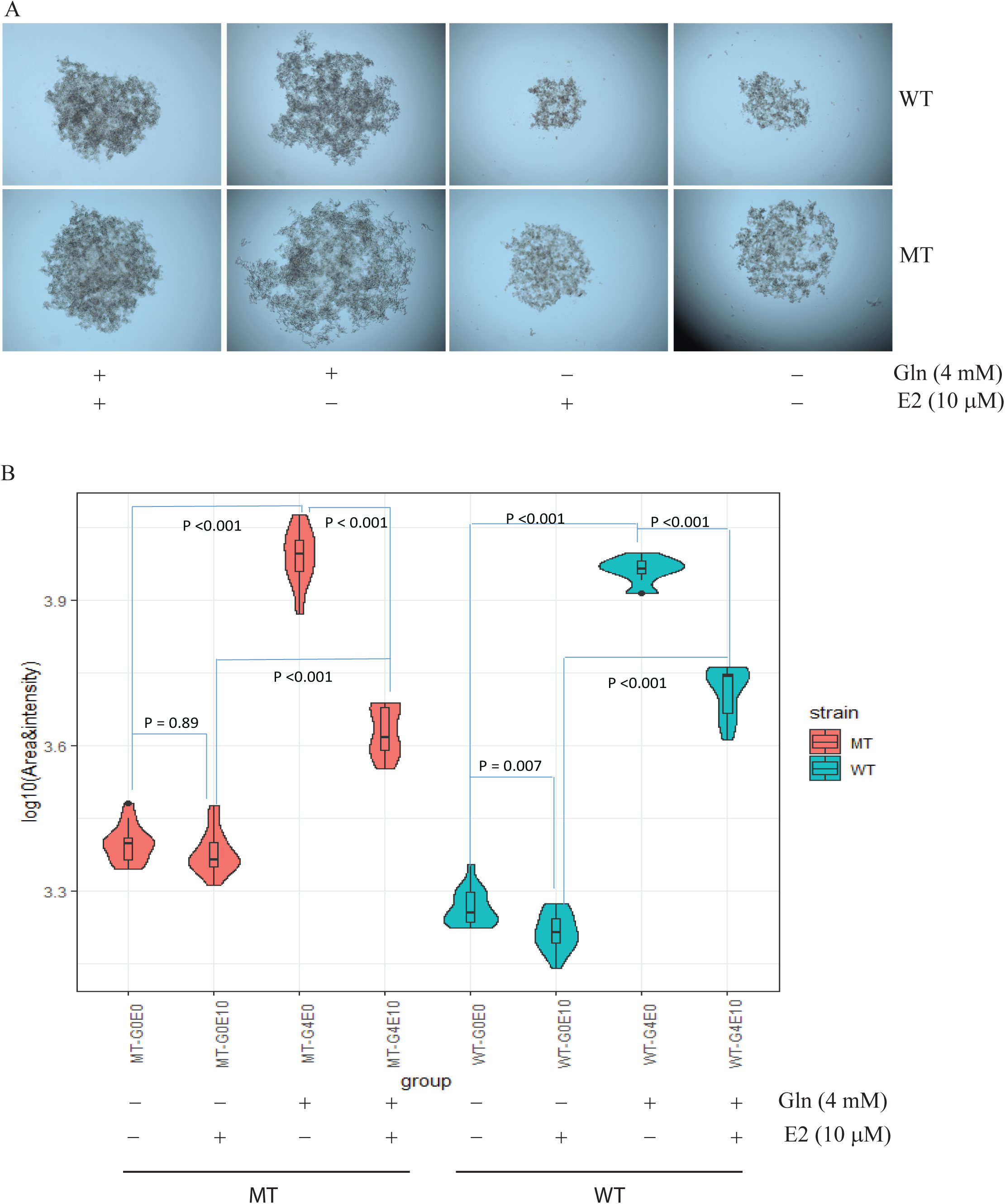
Effect of glutamine depletion and estradiol supplementation on 3D spheroid formation in vitro of human female CRC SW48 cells. (**A**) Representative images of 3D spheroids grown under glutamine deplete (0 mM) or replete (4 mM) DMEM medium with 10 μM or 0 μM estradiol. (**B**) Violin and box plots of 3D spheroid size and log intensity for SW48 *KRAS* MT and WT grown under different conditions. Post-hoc Tukey multiple comparison correction was applied after general linear model (GLM) for the ANOVA test. n = 12 each group per condition. Violin and box plots in cyan are for *KRAS* WT cells, and in Congo pink for the *KRAS* MT cells.

### Estradiol inhibits 3D spheroid formation in a nutrient-dependent manner in KRAS mutant and WT cells

To assess the effect of 17β-estradiol (E2) on cell proliferation, both SW48 *KRAS* MT and WT cells were grown in 4 mM glutamine replete DMEM, containing either 0 (0.1% DMSO as a negative control), or 10 µM E2 for 3D organoid assays. After 96-hours of incubation in glutamine replete DMEM, 10 µM E2 treatment significantly inhibited the 3D spheroid formation of both *KRAS* MT and WT compared to cells grown in the absence of E2 (p < 0.001) (**Figure 2**). The inhibition rate on average was 56.6% for the *KRAS* MT which was significantly higher than that for the *KRAS* WT cells of 43.3% (p = 1.05×10^-5^). When the cells were grown in 0 mM glutamine deplete DMEM, the inhibitory effect of E2 was only observed in the *KRAS* WT cells (p = 0.007), but not in the *KRAS* MT cells (p = 0.89). The inhibition rate of E2 was 10.9% for the *KRAS* WT, which was significantly higher than the rate of 4.3% for the *KRAS* MT on average under nutrient deplete conditions (p = 0.030). Therefore, E2 has a stronger inhibitory effect on *KRAS* WT cells compared to *KRAS* MT cells, and E2 has no inhibitory effect on *KRAS* MT cells in glutamine deplete conditions.

### Glutamine depletion upregulates ASNS in KRAS MT compared to WT cells, and estradiol supplementation has no effect

We previously reported that female CRC patients have a nutrient depletion metabolic phenotype characterized by increased ASNS-mediated asparagine production, and an unfavorable prognosis (14). To determine the effect of nutrient depletion and E2 on *ASNS* expression, we created an *in vitro* 3D CRC spheroid model using glutamine depletion as a proxy for nutrient depletion in the tumor microenvironment and supplemented these cells with E2 to determine the effect of this hormone on *ASNS*. Our results show that the expression of *ASNS* was significantly upregulated in *KRAS* MT compared to *KRAS* WT cells (log2 fold change = 1.22, FDR = 1.54 × 10^-4^) (**Figure 3A**). In nutrient replete conditions, *ASNS* was not altered between *KRAS* MT and WT CRC cells (log2 fold change of *ASNS* = 0.29 (FDR = 0.20). We then examined the effects of E2 supplementation on the cell lines. We observed that E2 supplementation had no additive effect on ASNS expression when comparing SW48 MT to WT cells (**Figure 3A**). When comparing the effect of E2 on MT and WT lines independently, we observed that E2 only had an effect by decreasing *ASNS* when comparing 0 μM versus 10 μM E2 supplementation in KRAS MT cells with nutrient replete conditions (log2 fold change = –0.75, FDR = 3.53×10^-4^) (**Figure 3B**).

**Figure 3.**
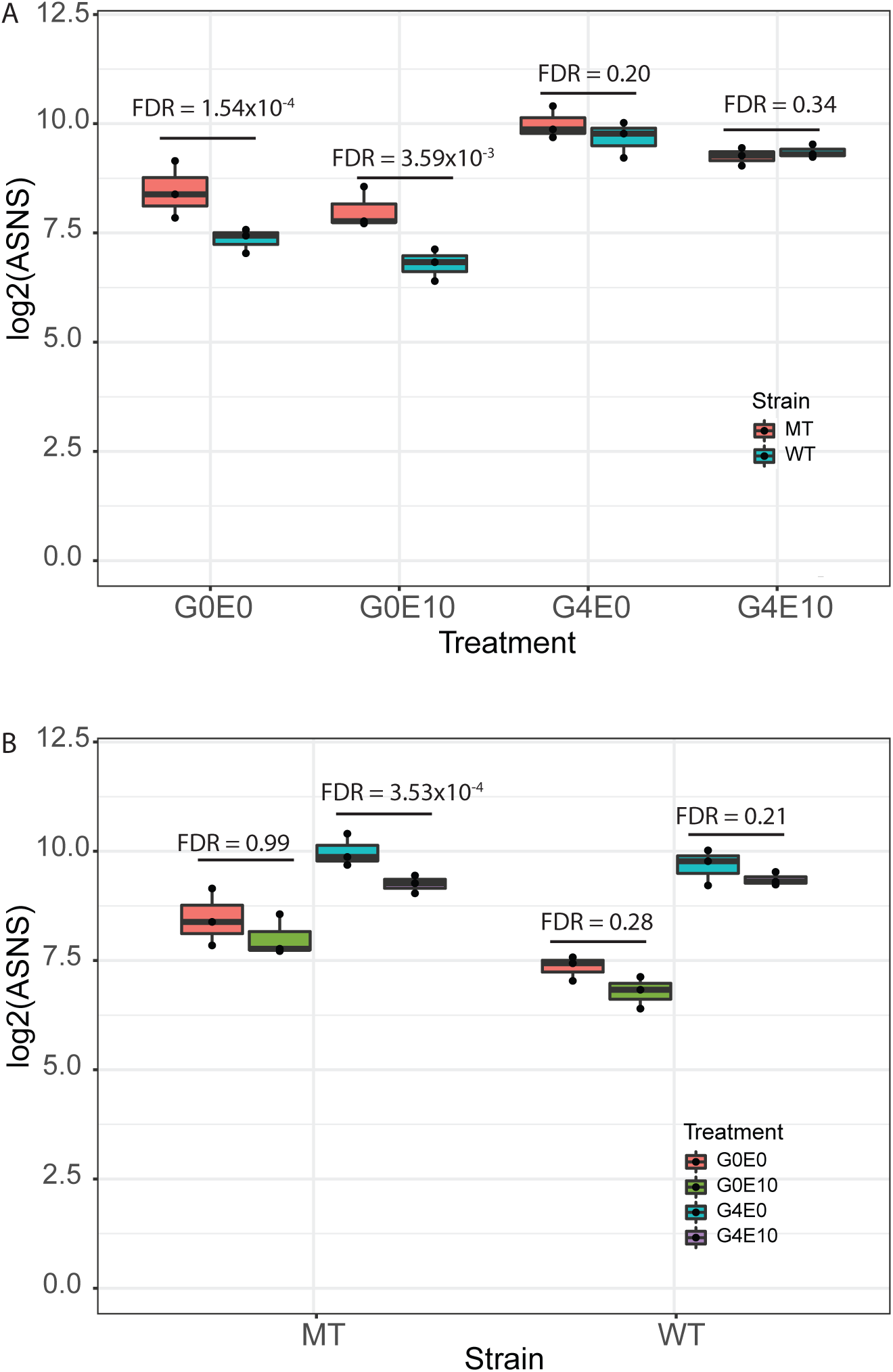
Effect of glutamine depletion and estradiol supplementation on the expression *ASNS* gene in in vitro 3D spheroid models. (**A**) Boxplot comparison of *ASNS* gene expression in log2 in MT vs. WT grown under different conditions without estradiol supplementation, or with 10 μM estradiol supplementation. **(B)** Boxplot comparison of *ASNS* gene expression in log2 between different culture conditions in MT and WT without estradiol supplementation, or with 10 μM estradiol supplementation.

Therefore, glutamine depletion increases *ASNS* expression in *KRAS* MT compared to *KRAS* WT cells, and E2 does not alter this effect. Furthermore, under glutamine replete conditions, 10 μM E2 supplementation decreases ASNS expression in *KRAS* MT cells but this effect is not observed in *KRAS* WT cells.

### Glutamine depletion upregulates GPER1 in KRAS MT compared to WT cells, and estradiol supplementation has no effect

As GPER1 can modulate PI3K/AKT signaling downstream of KRAS, we also examined *GPER1* expression in *KRAS* MT vs. WT cell lines after modulation of nutrient supply (i.e., glutamine) and E2 levels. Our results demonstrate that glutamine depletion significantly increases the expression of *GPER1* in *KRAS* MT compared to *KRAS* WT cells independent of E2 supplementation, similar to *ASNS*. The log2 fold change of *GPER1* expression was 0.78 at 0 μM E2 (FDR = 2.04×10^-4^) and 0.80 at 10 μM E2 (FDR = 3.96×10^-7^) (**Figure 4A**). Again, similar to the effects on *ASNS*, *GPER1* levels were not significantly altered between the *KRAS* MT and the WT cells under glutamine replete condition regardless of E2 supplementation. The log2 fold changes were 0.46 at 0 μM E2 (FDR = 0.22) and 0.45 at 10 μM E2 (FDR = 0.06) (**Figure 4A**). Further analyses of the effects of E2 on *GPER1* expression, showed no significant differences in expression within each cell line. Under glutamine deplete conditions comparing 0 μM to 10 μM E2, the log2 fold change = –0.08 (FDR = 0.99) for *KRAS* MT and –0.12 (FDR = 0.93) for the *KRAS* WT. Under glutamine replete conditions, the log2 fold change = 0.40 (FDR = 0.16) for *KRAS* MT and 0.40 (FDR = 0.27) for the *KRAS* WT cells (**Figure 4B**). Therefore, similar to the results observed for ASNS, glutamine depletion increases *GPER1* expression in *KRAS* MT compared to *KRAS* WT cells, and E2 does not alter this effect. However, E2 supplementation does not alter *GPER1* in *KRAS* MT under nutrient replete conditions, as was observed for *ASNS*.

**Figure 4.**
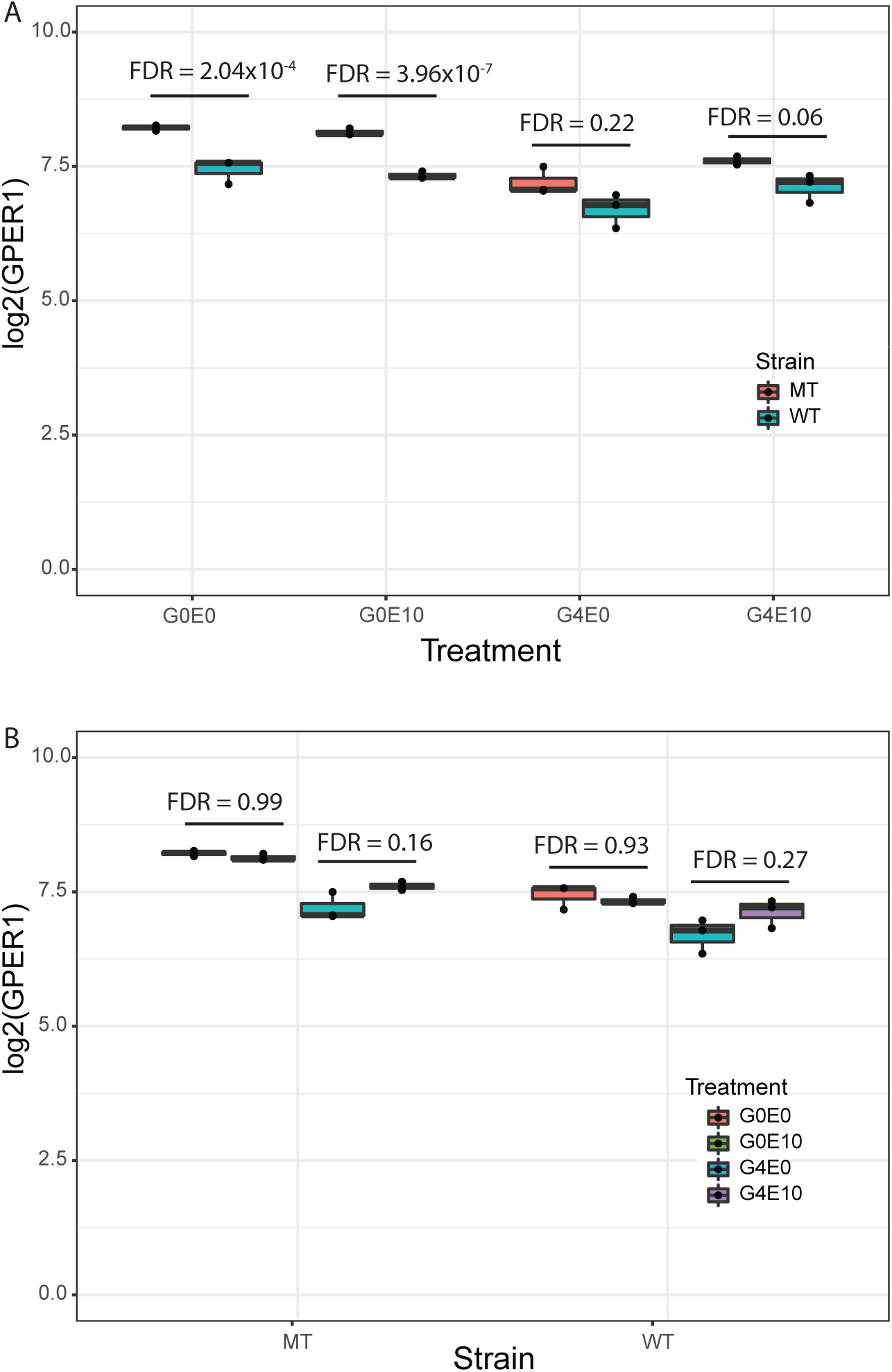
Effect of glutamine depletion and estradiol supplementation on the expression *GPER1* genes in in vitro 3D spheroid models. (**A**) Boxplot comparison of *GPER1* gene expression in log2 in MT vs. WT grown under different conditions. Upregulated expression of *GPER1* gene in the *KRAS* MT vs. WT cells when grown under glutamine depletion condition without estradiol supplementation, or with 10 μM estradiol supplementation. (**B**) Boxplot comparison of *GPER1* gene expression in log2 between different culture conditions in MT and WT with vs. without 10 μM estradiol supplementation.

### Glutamine depletion and estradiol supplementation have no effect on ESR1 and ESR2 in KRAS MT compared to WT cells

Given that GPER1 is an estrogen receptor, we examined whether other estrogen receptors were altered in these conditions. We observed that the nuclear estrogen receptors ERα (*ESR1*) and ERβ (*ESR2*) were not altered in their expression between *KRAS* MT and the WT cells, and E2 supplementation had no effect. *ESR1* expression was either not detectable or had very low counts in both the *KRAS* MT and WT cells; it is known that *ESR1* is not typically expressed in colon cancer (32). Under glutamine deplete conditions the log2 fold changes for *ESR2* were 0.09 (FDR = 0.93) without E2, and –0.10 (FDR = 0.88) with E2. Under glutamine replete conditions, the log2 fold changes for *ESR2* were 0.28 (FDR = 0.63) without E2 and –0.06 (FDR = 0.92) with E2 (**Figure S1A**).

### Pathway analysis

RNA-Seq was performed on KRAS MT and KRAS WT cells that were grown in nutrient deplete and replete conditions, with E2 treatment. **Figure S2A** shows a global alteration of transcriptome profiles between SW48 *KRAS* MT and WT CRC cell lines under glutamine deplete conditions prior to supplementation with E2. **Figure S2B** shows the alteration of transcriptome profiles between SW48 *KRAS* MT and WT CRC cell lines under nutrient deplete conditions. However, the number of significantly downregulated genes was less under the deplete conditions (796 vs. 1193, with 413 shared downregulated genes) as compared to the nutrient replete condition (based on 2-fold change and FDR<0.05), whereas the number of significantly upregulated genes was more (616 vs. 466, with 242 shared upregulated genes) (**Figure S2C**) (p<0.0001). Similarly, a global change in transcriptome profiles was observed in the glutamine deplete or replete conditions with 10 μM estradiol supplementation when comparing *KRAS* MT with WT cells (**Figures S2D–E**). The number of significantly downregulated genes was less under the deplete conditions in the presence of estradiol (890 vs. 1075, with 403 shared downregulated genes) as compared to the nutrient replete condition in the presence of estradiol (based on 2-folds change and FDR<0.05), whereas the number of significantly upregulated genes was more (750 vs. 540, with 230 shared upregulated genes) (**Figure S2F**) (p < 0.0001).

Ingenuity pathway analysis (IPA) was then used to identify pathways altered between KRAS MT and KRAS WT cells by treatment condition. The analysis shows that genes involved in cell cycle, cell adherence, cancer and inflammation were differentially expressed in *KRAS* MT vs. WT cells when grown under glutamine deplete condition without E2 supplementation. The six canonical pathways with a Z-score ≥2 (i.e., activated pathways) included SPINK1 pancreatic cancer, cyclins and cell cycle regulation, estrogen-mediated S-phase entry, cell cycle control of chromosomal replication, mitotic role of polo-like kinase and kinetochore metaphase signaling. The six pathways with a Z-score ≤-2 (i.e., inhibited pathways) included cell G1/S checkpoint regulation, senescence pathway, epithelial adherens junction signaling, MSP-RON signaling in macrophages and cancer cell pathways (**Figure 5A**). Similarly, the differentially expressed genes in *KRAS* MT vs. WT under glutamine deplete condition with E2 supplementation were involved in cell cycle, cancer and inflammation, which included three activated pathways of SPINK1 pancreatic cancer, cell cycle control of chromosomal replication and estrogen-mediated S-phase entry, and the six inhibited pathways of G1/S checkpoint regulation, role of CHK proteins in cell cycle checkpoint control, intrinsic prothrombin activation, MSP-RON signaling in macrophage and cancer cells, and xenobiotic metabolism PXR signaling (**Figure 5B**).

**Figure 5.**
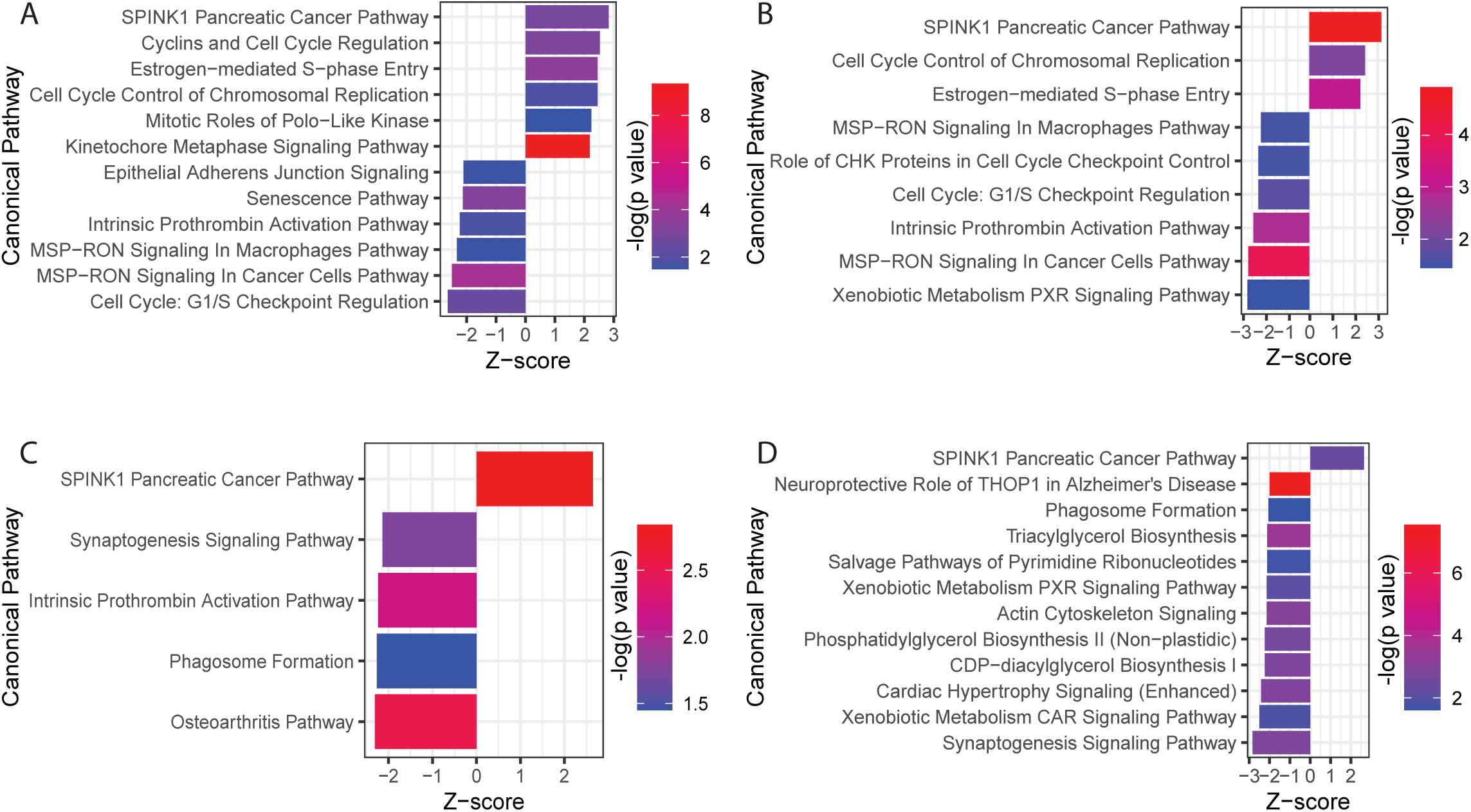
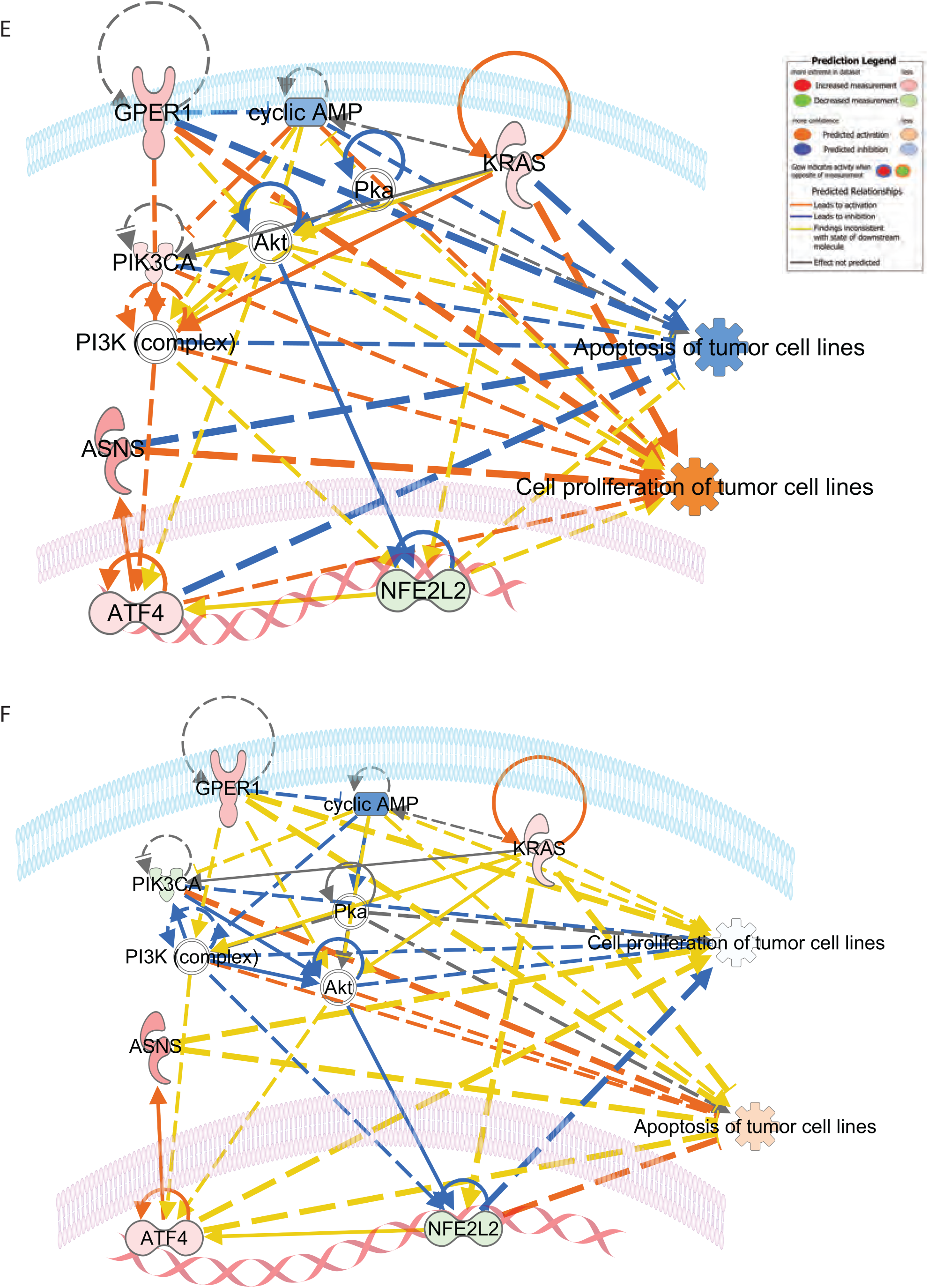
Ingenuity pathway analysis for differentially expressed genes (DEG) in KRAS MT vs. WT under different conditions. (**A**). Bar plots of the top canonical pathways for the DEGs under glutamine deplete condition without E2 supplementation, or (**B**). with E2 supplementation, and (**C**).glutamine replete condition without E2 supplementation, (**D**) or with E2 supplementation. (**E**). The network of KRAS, GPER1 and ASNS genes involved in the proliferation and apoptosis of tumor cells under glutamine deplete condition without E2 supplementation, and (**F**) with E2 supplementation. Orange line: leads to activation; blue line: leads to inhibition; yellow line: findings inconsistent with state of downstream molecule; grey line: effect not predicted. Molecules with pink: increased; Molecules with green: decreased. Functions with orange: predicted activation, functions with blue: predicted inhibition. Line with arrow: activation; line with stop sign: inhibition

When the cells were grown under glutamine replete conditions without E2 supplementation, enriched pathways in *KRAS* MT vs. WT included activation of SPINK1 pancreatic cancer pathway, and inhibition of four pathways of synaptogenesis signaling, phagosome formation, osteoarthritis, and intrinsic prothrombin activation pathways (**Figure 5C**). Again, when the cells were grown under glutamine replete condition with E2 supplementation, only activated SPINK1 pancreatic cancer pathway was enriched. However, there were 11 inhibited pathways included synaptogenesis signaling, xenobiotic metabolism CAR signaling, cardiac hypertrophy signaling, CDP-diacylglycerol biosynthesis I, phosphatidylglycerol biosynthesis II, actin cytoskeleton signaling, xenobiotic metabolism PXR signaling, salvage pathways of pyrimidine ribonucleotides, triacylglycerol biosynthesis, phagosome formation, and neuroprotective role of THOP1 in Alzheimer’s disease (**Figure 5D**). Thus, glutamine depletion leads to the inhibition of MSP-RON signaling in macrophage and cancer cells and G1/S checkpoint regulation, whereas glutamine repletion results in the inhibition of phagosome formation and synaptogenesis signaling in *KRAS* MT compared to *KRAS* WT cells. E2 supplementation leads to the inhibition of xenobiotic metabolism PXR signaling in in *KRAS* MT compared to *KRAS* WT cells under both glutamine deplete and replete conditions. The activation of SPINK1 pancreatic cancer pathway is shared for all four culture conditions in *KRAS* MT compared to *KRAS* WT cells.

Further network analysis using IPA shows a potential link between these pathways and GPER and ASNS. Increased *GPER1* and *ASNS* expression in KRAS MT compared to KRAS WT cells grown in glutamine deplete conditions without E2 supplementation were linked to increased cell proliferation and decreased apoptosis of tumor cells (**Figure 5E**), whereas with E2 supplementation, the opposite is predicted with the activation of apoptosis and shows a protective effective of E2 in CRC (**Figure 5F**). However, upregulated *GPER1*, and *ASNS* inhibited the E2-induced apoptosis in *KRAS* MT cells when under the depleted glutamine conditions supplemented with E2 (**Figure 5F**). This appears to be mediated through the differential effects of ASNS and GPER, the downregulation of PIK3CA, inhibition of protein kinase A (PKA) and nuclear factor erythroid 2-related factor (NFE2L2) (**Figure 5F**).

### The association of high GPER1 and ASNS expression with poorer survival is sex-and stage-dependent in patients with colon cancer

Patients with advanced stage CRC demonstrate significantly lower serum glutamine concentrations compared to those with the early-stage disease, suggesting that nutrient depletion exists in patients with advanced compared to early-stage CRC (33, 34). Moreover, overexpression of HIF1α in CRC at advanced-stage CRC compared to early-stage CRC suggests a more hypoxic tumor microenvironment in the former (35, 36). The association between *GPER1* and *ASNS* expression levels and the survival of patients with primary CRC was assessed in a GDC TCGA colon cancer dataset using Kaplan-Meier survival analysis stratified by sex and disease stage, under the hypothesis that advanced stage tumors are more nutrient deplete. We observed that the association between *GPER1* expression and overall survival was cancer stage-dependent in females, but not in males (**Figure 6A–D**). A significant and positive association was found between high *GPER1* expression and poor overall survival in females with advanced stage colon cancer (III-IV) (log-rank p = 0.034, **Figure 6A**). The 3-year survival rates were 46.0% (95% CI: 27.0-78.3%) for female patients with a high *GPER1* expression level, and 70.5% (95% CI: 57.2-87.0%) for those with low *GPER1*. However, an opposite association was observed in females with early-stage disease (I-II); high *GPER1* expression levels had a superior overall survival than those with a low expression, and the association was borderline but not statistically significant (log-rank p = 0.063) (**Figure 6B**). The 3-year survival rates were 100% for females with a high *GPER1* expression level, and 88.4% (95% CI: 80.4-97.2%) for those with low expression. No significant association was found between *GPER1* expression and overall survival in males with a disease at advanced stage (log-rank p = 0.96) (**Figure 6C**). The 3-year survival rates were 71.8% (95% CI: 55.9-92.2%) for males with a high *GPER1* expression level, and 56.8% (95% CI: 42.9-75.3%) for males with low expression. Similarly, no significant association was found in males with a disease at early stage (log-rank p = 0.35) (**Figure 6D**). The 3-year survival rates were 92.3% (95% CI: 82.6-100%) for males with a high *GPER1* expression level, and 89.1% (95% CI: 82.0-96.8%) for those with a low expression.

**Figure 6.**
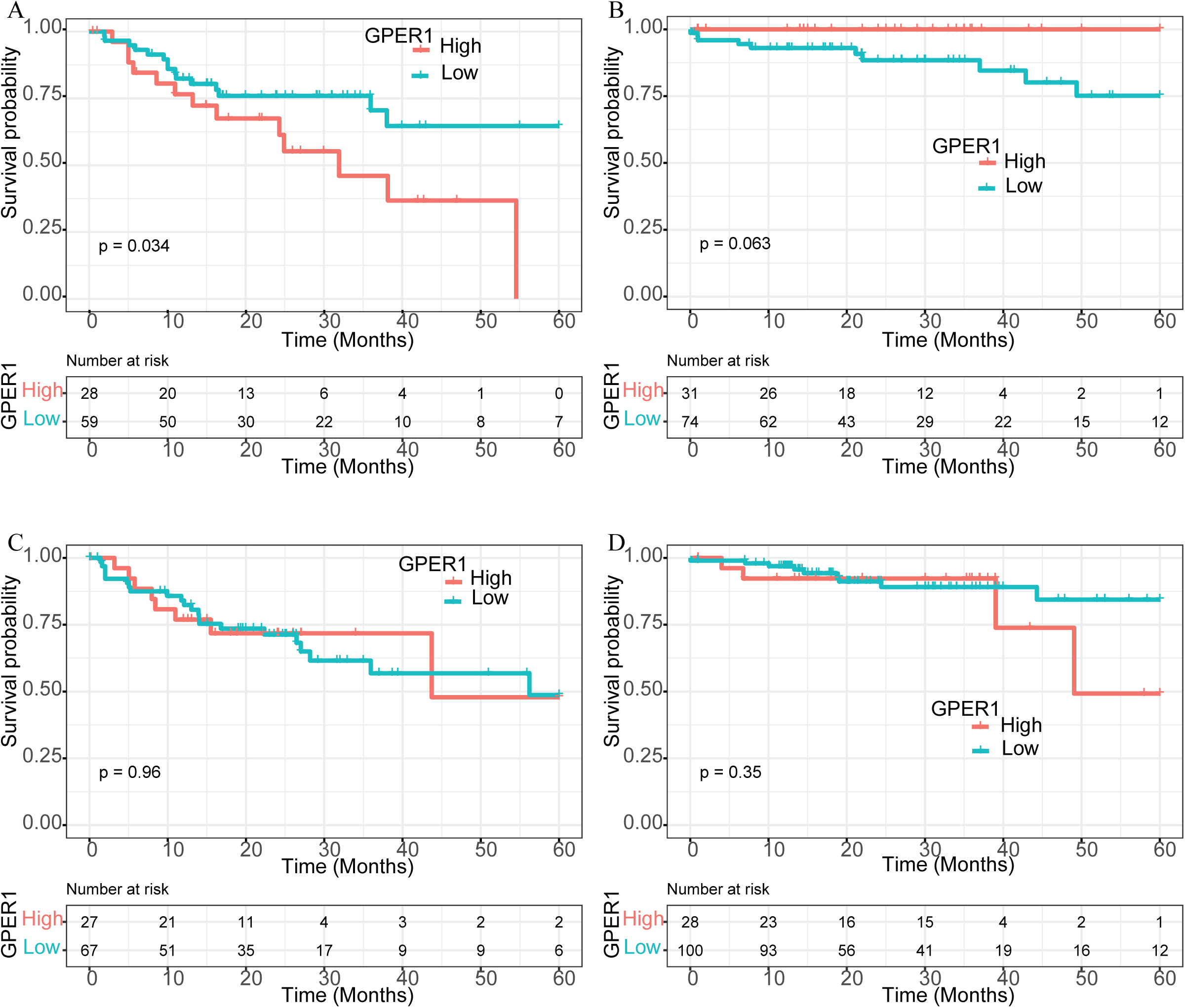
Kaplan-Meier overall survival stratified by *GPER1* expression levels in a TCGA CRC dataset. (**A**) female patients at advanced stage, **(B)** female patients at early stage, (**C**) male patients at advanced stage, and (**D**) male patients at early stage

Consistent with these results, the association of *ASNS* expression and patient overall survival was sex-and stage-dependent. High expression of *ASNS* was significantly associated with poorer survival in females with a disease at advanced stage (log-rank p = 0.048) (**Figure 7A**). The 3-year survival rates were 48.9% (95% CI: 31.6-75.6%) for female patients at advanced stage with a high *ASNS* expression level, and 75.2% (95% CI: 63.4-89.2%) for females at advanced stage with a low expression. No significant associations were found in females with an early-stage disease (log-rank p = 0.71) (**Figure 7B**). The 3-year survival rates were 95.6% (95% CI: 89.7-100%) for females at early stage with a high *ASNS* expression, 88.7% (95% CI: 79.4-99.0%) for females at early stage with low expression. Again, no significant association was found between *ASNS* and overall survival in males with either advanced (log-rank p = 0.63, **Figure 7C**) or early-stage disease (log-rank p = 0.56, **Figure 7D**). For males with advanced stage disease, the 3-year survival rates were 64.8% (95% CI: 51.0-82.4%) for those with a high *ASNS*, and 51.7% (95% CI: 33.1-80.8%) for those with a low *ASNS*. In contrast, for males with early-stage disease, the 3-year survival rates were 90.1% (95% CI: 82.8-98.0%) for those with a high *ASNS*, and 89.4% (95% CI: 79.5-100%) for those with a low *ASNS*. Therefore, high *GPER1* and *ASNS* expression associates with poorer survival for female patients with advanced disease stage only when the tumor is more nutrient deplete.

**Figure 7.**
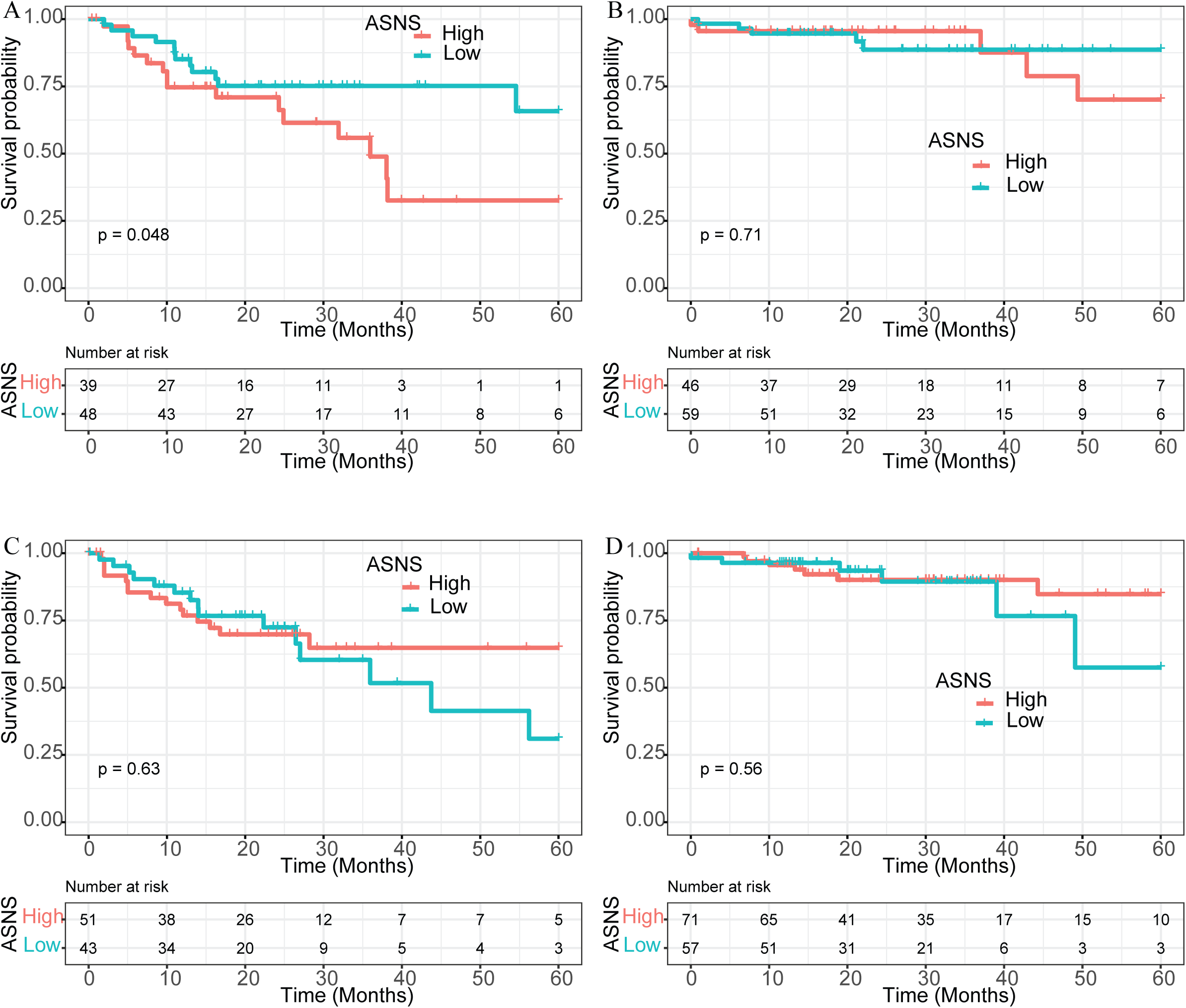
Kaplan-Meier overall survival stratified by *ASNS* expression levels in a TCGA CRC dataset. (**A**) female patients at advanced stage, (**B**) female patients at early stage, (**C**) male patients at advanced stage, and (**D**) male patients at early stage.

## Discussion

In this study, we demonstrate the impact of nutrient depletion and the sex-steroid hormone E2 on tumor cell proliferation, and the expression of *ASNS* and *GPER1* in CRC. Using a 3D spheroid model, we found a significant reduction in the size of 3D spheroid in both *KRAS* MT and WT cells grown in glutamine deplete medium (0 mM) compared to glutamine replete medium (4 mM). E2 supplementation significantly inhibited the proliferation of both cell types when grown in the glutamine replete medium. However, E2 supplementation had no effect on the *KRAS* MT cells when grown in the glutamine deplete medium, but significantly inhibited the WT cells. These results suggest that CRC growth depends on tumor nutrient availability, *KRAS* mutation status and E2; E2 exposure decreases CRC cell growth but has no significant effect when the cells harbor *KRAS* mutations and are cultured under nutrient deplete conditions.

These findings support observations from previous studies that sex-steroid hormones can modulate tumor growth. Studies in mouse models have suggested that the endogenous female hormone estrogen has a protective role against the development of colon cancer, and exogenous estradiol replacement could protect ovariectomized female mice from azoxymethane (AOM)/dextran sulfate sodium (DSS)-induced colitis and colon carcinogenesis (37). It has also been shown that female patients with CRC, and patients younger than 65 years old have a survival advantage over age-matched males regardless of disease stage at diagnosis (38). In addition to the protective role of endogenous estrogen hormones, the use of oral contraceptives, and postmenopausal HRT also protect women from CRC development (17, 39, 40). A meta-analysis also indicated that combined estrogen-progestogen therapy reduces the risk of CRC by 26% (RR = 0.74, 95% CI: 0.68–0.81) and estrogen-only therapy reduces the risk of CRC by 21% (RR = 0.79, 95% CI: 0.69–0.91) (17). A postmenopausal cohort study in Norway showed that current menopausal HRT use decreased CRC risk (39). In addition, the Women’s Health Initiative trial of estrogen plus progestin in postmenopausal women demonstrated a 44% reduction of risk of invasive CRC in the hormone supplemented group vs. placebo group (HR = 0.56, 95% CI: 0.38-0.81) (40).

In our data, as expected, glutamine depletion significantly inhibited tumor cell growth in both *KRAS* MT and WT cells. Although glutamine is a non-essential amino acid, it is a substrate for asparagine, and glutathione which neutralizes reactive oxygen species (ROS). Glutamine also contributes to the generation of energy and substrates for macromolecule synthesis through glutaminolysis and by entering the tricarboxylic acid (TCA) cycle (25). Glutaminolysis provides nitrogenous and carbon compounds for the synthesis of nucleotides, lipid, and non-essential amino acids to sustain the demand for rapid proliferation and growth of tumor cells. Furthermore, although glutamine depletion leads to the inhibition of tumor growth, asparagine supplementation can rescue glutamine depletion-induced suppression of tumor cell growth and viability (4). In response to glutamine depletion, *ASNS* is upregulated to enhance intracellular asparagine biosynthesis, particularly in *KRAS* mutant CRC cells (4), in which activated oncogenic *KRAS* signaling leads to the increase of ATF4 via PI3K/AKT signaling (7). As an upstream regulator, ATF4 further promotes the expression of *ASNS*, consequently making the cells adaptive to glutamine depletion-induced apoptosis (4, 7). Moreover, *ASNS* knockout melanoma cells were shown to be sensitive to the depletion of non-essential amino acids in vitro, with a rapid decrease in cellular proliferation and cellular asparagine levels which increase aspartate levels (the substrate of asparagine) (41). In line with these previous findings, we observed that *ASNS* expression was upregulated in *KRAS* MT vs. WT cells when grown in glutamine-deplete medium regardless of E2 supplementation. Moreover, in the TCGA colon cancer dataset we also found that high *ASNS* expression significantly associated with poor overall survival in females at advanced stage, but not in early stage or in male patients. Given that ASNS plays an important role in tumor growth and metastasis in human cancer (10, 41–43), the finding of upregulated *ASNS* expression in this study also suggests one of the mechanisms by which *KRAS* MT CRC is more aggressive than *KRAS* WT CRC. Since the sample size is relatively small in the subgroups, we could not further analyze the association between *ASNS* expression and patient survival based on *KRAS* status.

ERα and ERβ are two nuclear receptors that regulate widespread gene expression by engaging with the ligand estrogen. ERα and ERβ, however, have distinct functions in tumor development and progression, with ERα being a tumor promoter, and ERβ protective. *ERS1* has limited or no expression in the colon tissue, and *ESR2* is the predominant estrogen receptor (ER) expressed in both normal and malignant colonic epithelium, although *ESR2* expression is reduced during colonic tumorigenesis relative to normal colon tissues (32). In agreement with these results, we found undetectable or very low levels of *ESR1* in the CRC cells used in our study. There was no significant difference in *ESR2* expression between the two cell lines, and E2 supplementation had no effect on *ESR2* expression. Interestingly, we observed that the expression of *GPER1*, a transmembrane bound estrogen receptor, was upregulated in *KRAS* MT cells compared to WT cells under glutamine depletion. We also observed that under glutamine deplete conditions, E2 supplementation had no effect on the growth of *KRAS* MT cells but inhibited the growth of *KRAS* WT cells.

In the TCGA colon cancer dataset, we found that *GPER1* high (vs. low) expression positively associates with worse survival for female patients with CRC at advanced stage, and an opposite direction was observed for the association for females at early-stage CRC. However, no significant association existed between *GPER1* expression and overall survival in males with either early or advanced stage. This underscores the importance of the tumor microenvironment and nutrient availability for growing tumors and their interplay with important signaling molecules in. sex-specific manner. Advanced stage tumors are more hypoxic, have restricted blood supply and nutrients. This data shows that the potential actions of GPER1 in females is dependent on the tumor microenvironment but has no effect in males. These results are in accordance with a previous study showing high *GPER1* expression was associated with poor survival exclusively in women (19). Furthermore, the GPER agonist G-1, suppressed *in vivo* cell growth and promoted apoptosis in CRC cell lines of HCT116 and SW480 (26). In contrast, estrogen-driven GPER1 activation accelerated the proliferation of CRC cell lines in another study (27). In our study, we used a 3D spheroid model to grow both *KRAS* MT and WT cells, which mimics the insufficient nutrient and hypoxia microenvironment of tumor mass *in vivo*. Increased *GPER1* expression was observed in *KRAS* MT compared to WT cells under glutamine deplete conditions regardless of E2 supplementation. In combination with the observation of the inhibition of cell growth with E2 supplementation on *KRAS* WT cells but not in *KRAS* MT cells in nutrient deplete conditions, it is possible that GPER1 upregulation could promote tumor cell survival under glutamine deplete conditions when supplemented with E2 in *KRAS* MT cells. Pathway analysis shows that under nutrient deplete conditions, upregulated GPER1 inhibits the E2-induced apoptosis in *KRAS* MT cells and higher than it does for *KRAS* WT (**Figure 5F**). This is supported by the observation of sustained cell growth of *KRAS* MT greater than the WT cells in our 3D organoid model under the deplete glutamate conditions after E2 supplementation. We hypothesize that in *KRAS* MT cells there is a coordinated effect on these two processes (cell proliferation and apoptosis) by ATF4, ASNS, PIK3CA, PKA and GPER1 as outline in **Figure 5E** and **5F**, which have an opposing effect in *KRAS* WT cells.

Thus, it is possible that nutrient depletion induces the upregulation of important metabolic drivers that increase tumor growth when normal nutrient supply is prohibited (e.g., under times of hypoxia). This appears to be particularly important for female *KRAS* MT CRCs wherein both ASNS-driven increases in asparagine and procarcinogenic signaling via GPER increase tumor growth and associate with poorer prognosis. GPER1 is linked to *ASNS* regulation via its regulatory effects on PI3K/AKT through cyclic adenosine monophosphate (cAMP) (23–25). Upregulated *GPER1* and *ASNS* genes in *KRAS* MT vs. WT when they were grown under glutamine deplete conditions could help to tumor cell proliferation and inhibit apoptosis.

The tumor microenvironment is of great importance due to its role in providing adequate nutrient supply to sustain growth. Our study utilizes a 3D organoid model and showed that when nutrient supply is limited, metabolic rescue mechanisms such as *ASNS*-driven amino acid production increase, but only in *KRAS* MT CRC cells, a type of CRC that is more difficult to treat. Notably, GPER1, which is nutrient sensitive, was also increased in *KRAS* MT cells in these nutrient poor conditions. Interestingly, although GPER1 is an estrogen receptor, and *GPER1* and *ASNS* associate with poorer survival in advanced stage female patients (but not males), supplementation with E2 had no additional effect on *GPER* and *ASNS* expression. We also show that E2 has no effect on *KRAS* MT cell growth when nutrient supply is limited; however, E2 decreases the growth of *KRAS* WT cells. These results suggest that GPER and ASNS are critical rescuers of tumor growth in *KRAS* MT CRC when the nutrient supply in the colon tumor microenvironment is limited.

To our knowledge, this is the first study to provide fundamental knowledge on *ASNS* and *GPER1* expression in the link of nutrient supply and *KRAS* mutations in CRC, shedding additional light on the mechanisms behind sex differences in metabolism and growth in CRC. Moreover, ASNS and GPER1 are potential therapeutic targets that can be exploited to manage and control CRC, and the selective inhibition of glutamine metabolism in tumor cells may improve immunotherapy in patients with *KRAS* mutants.

## Acknowledgements

This work was funded by American Cancer Society Research Scholar Grant 134273-RSG-20-065-01-TBE (CJ). This publication was also made possible by CTSA Grant Number UL1 TR001863 from the National Center for Advancing Translational Science (NCATS), components of the National Institutes of Health (NIH), and NIH roadmap for Medical Research, as well as NCI R37CA259363 (JR)

## Supplementary Figure legends

**Figure S1.**
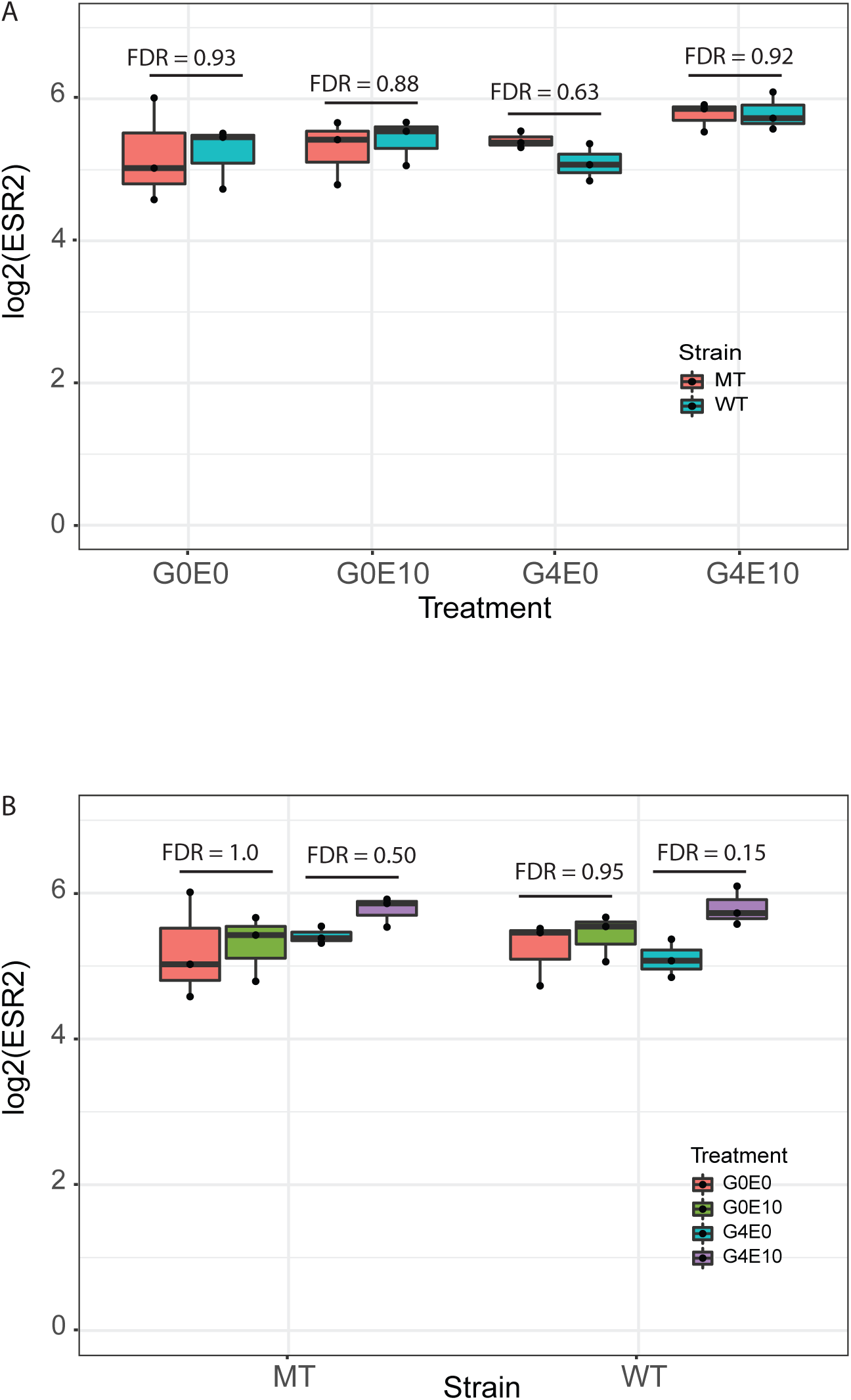
Effect of glutamine depletion and estradiol supplementation on *ESR2* expression. (**A**) No significant differences were observed in *ESR2* expression level between the *KRAS* MT vs. WT grown under glutamine depletion or repletion in the absence or presence of 10 μM estradiol. (**B**) or between the absence and presence of 10 μM estradiol.

**Figure S2.**
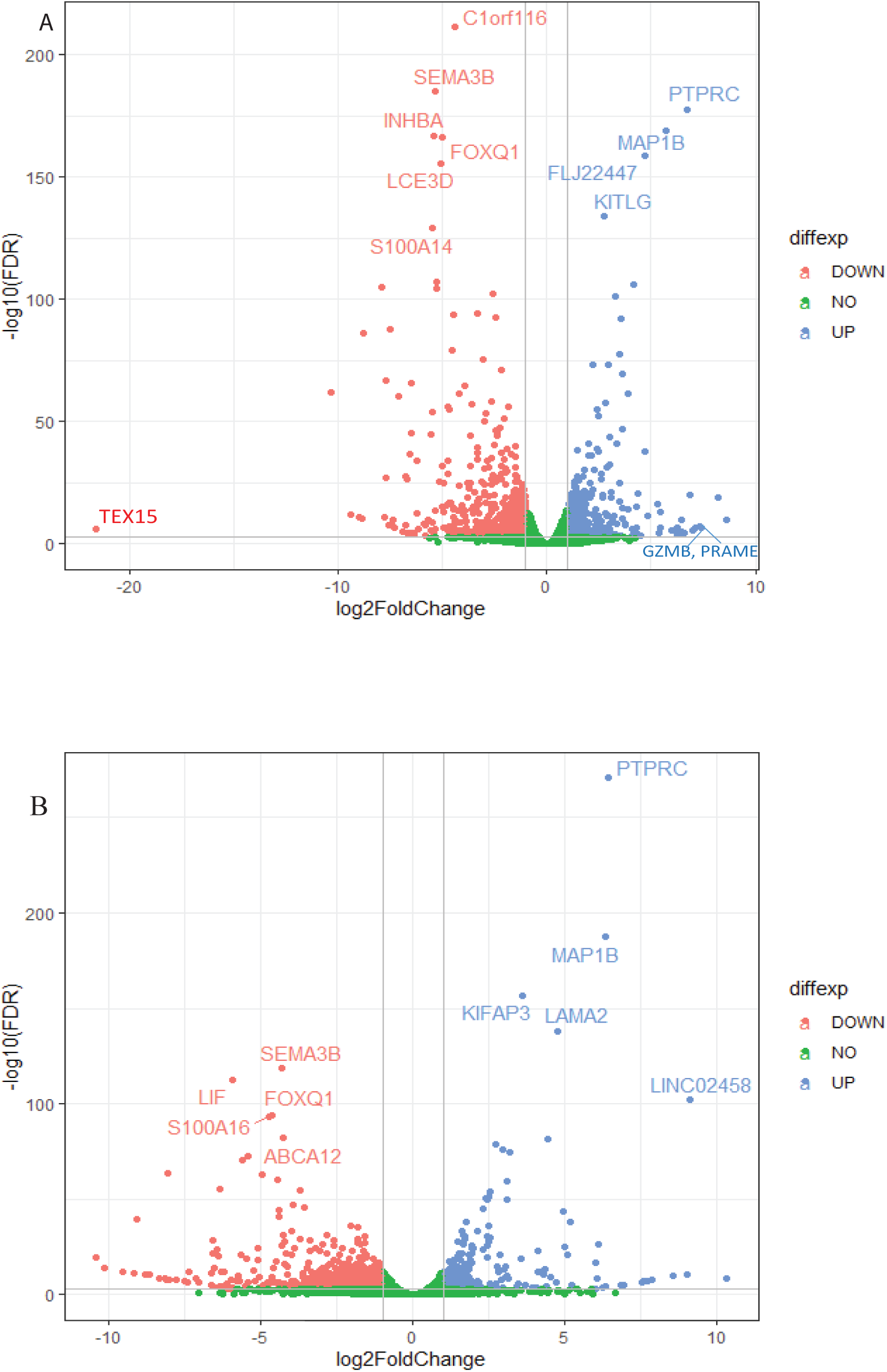
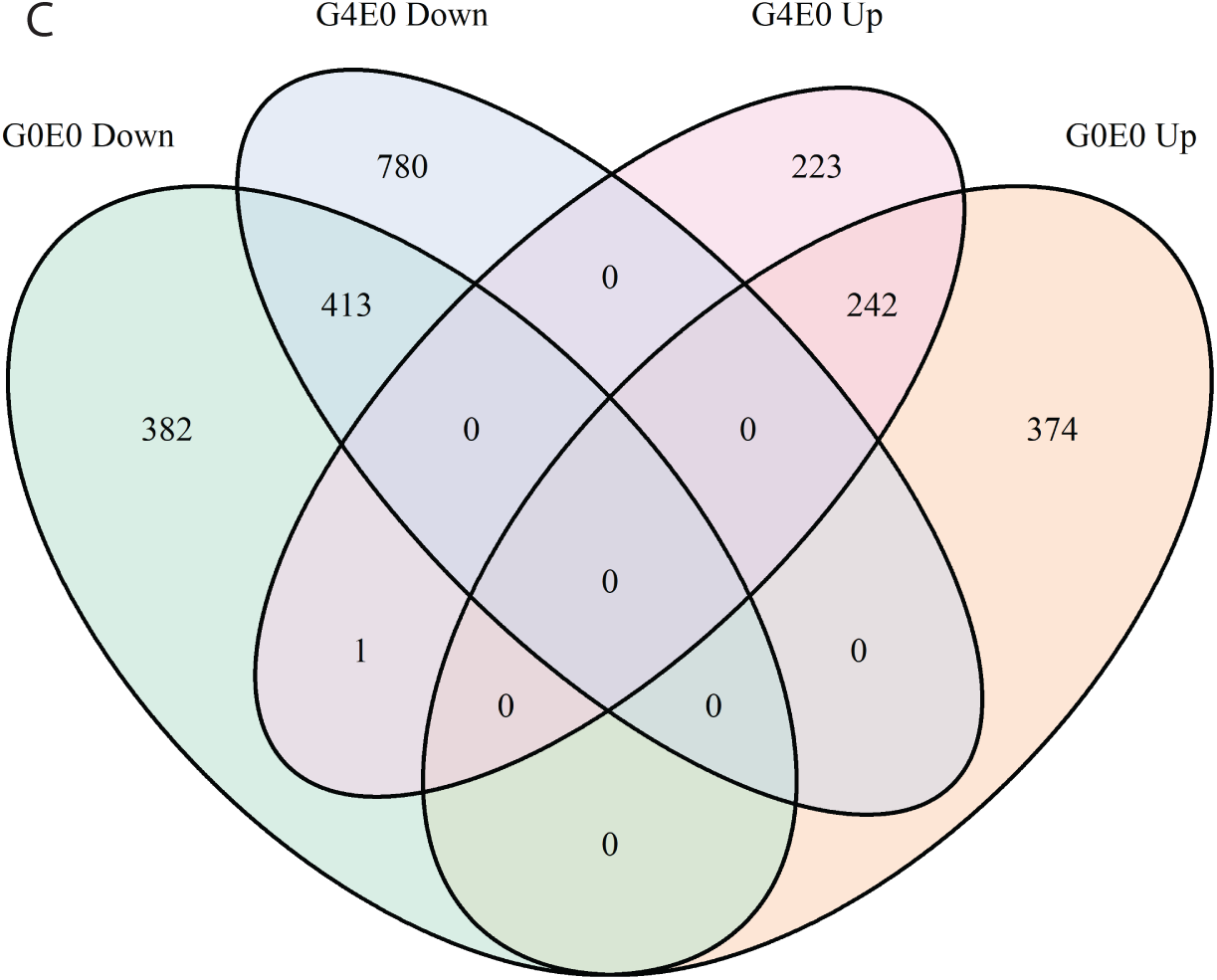
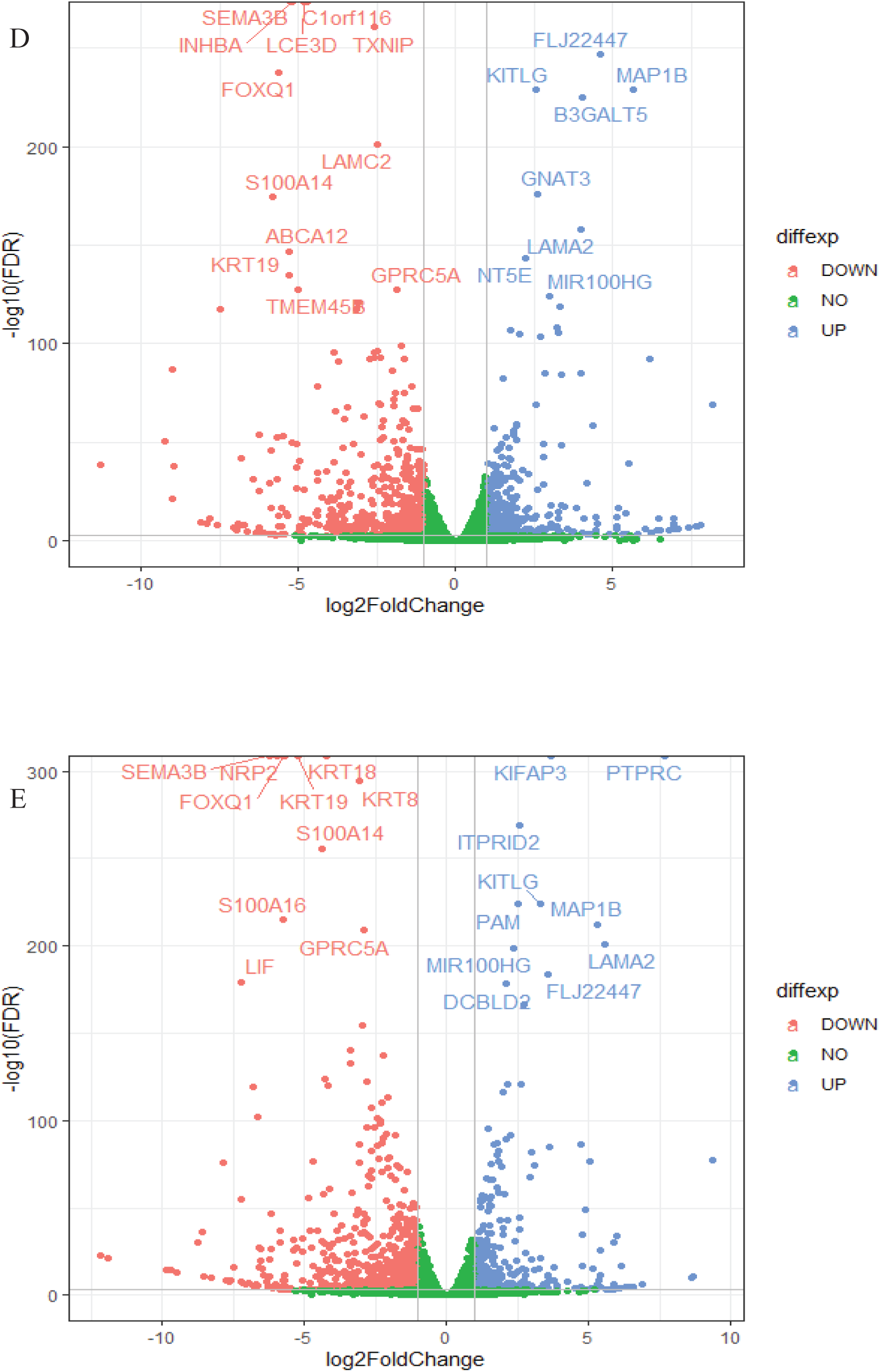
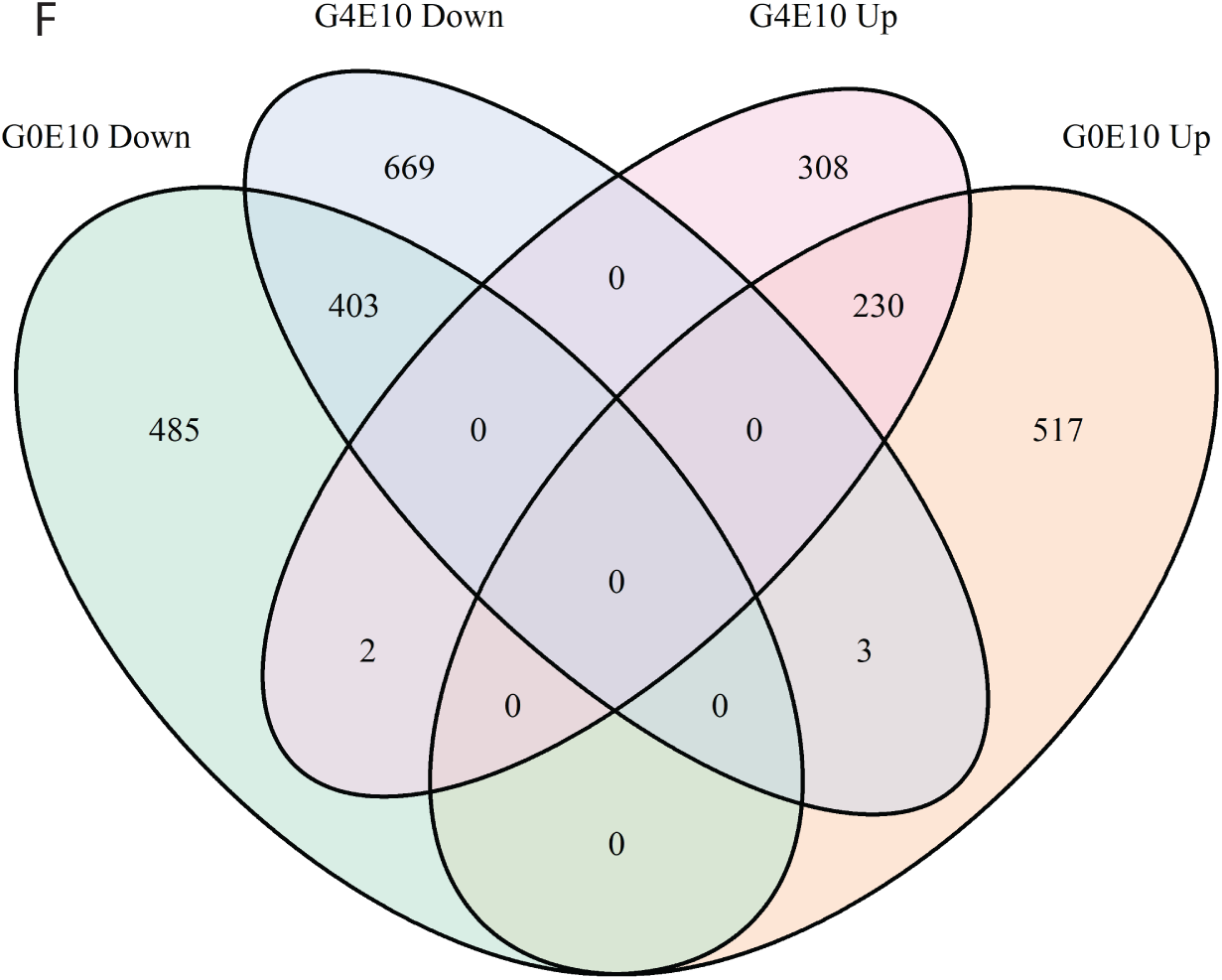
Volcano plots and Venn diagram of transcriptomic profiling in human female CRC SW48 *KRAS* MT and WT cells in in vitro 3D spheroid model. (**A**) Globally differential transcriptomic expression in *KRAS* MT vs. WT cells when grown under glutamine deplete, (**B**) or under glutamine repletion (4 mM) condition without estradiol supplementation, (**C**) Venn diagram of down-or up-regulated genes for cells under glutamine deplete (0 mM) or repletion (4 mM) condition without estradiol supplementation; (**D**) Globally differential transcriptomic expression in *KRAS* MT vs. WT cells when grown under glutamine depletion (0 mM) condition with 10 μM estradiol supplementation, (**E**) or under glutamine repletion with 10 μM estradiol supplementation. (**F**) Venn diagram of down-or up-regulated genes for cells under glutamine deplete (0 mM) or repletion (4 mM) condition with 10 μM estradiol supplementation.

